# Disruption of the insulin signaling pathway in *C. elegans* dramatically increases male longevity and enhances reproductive health late in life

**DOI:** 10.1101/2025.11.04.686639

**Authors:** Rose S. Al-Saadi, Hannah B. Lewack, Patrick C. Phillips

**Author notes:** Corresponding author Correspondence to Patrick Phillips.

## Abstract

Males and females are known to have dramatically different health and lifespan trajectories, but the underlying basis for these differences is only now being fully investigated^1^. In the *Caenorhabditis elegans* nematode model system, most aging studies have been conducted with hermaphrodites, and little is known about male-specific responses to pro-longevity mutations. Several previous studies have used the auxin-inducible degron system to degrade the insulin-like DAF-2/IGF-1 receptor in hermaphrodites, finding that both ubiquitous and tissue-specific degradation can extend lifespan^2–4^. Here we show that ubiquitous degradation of DAF-2 in male *C. elegans* increases median lifespan by more than 440%, one of the longest lifespan extensions by a single intervention to date. Conversely, degrading DAF-2 in the male germline decreased lifespan, opposite of its effect in hermaphrodites^3^. Using male mating and reproductive success as a meaningful ecological and neurophysiological measure of healthspan, we found that ubiquitous degradation of DAF-2 greatly prolongs reproductive health, likely by prolonging function of the male intromittent organ in the tail. This work highlights the importance of studying sex differences in aging and highlights the utility of using *C. elegans* males to understand the underlying basis of enhanced lifespan and healthspan.

## Main

Sex underlies pervasive differences in aging and longevity between males and females^5^. Such differences are seen in human populations, as well as most other mammalian species^6,7^. While complex social structures and other environmental factors undoubtedly contribute to this phenomenon, sex-based differences in longevity are nevertheless observed in social groups in which both sexes share similar lifestyles^8,9^. Precisely how sex regulates aging and longevity, especially at a molecular level, is still largely understudied. This is in part due to the complex nature of aging, and in part due to limited emphasis in research programs on how sex modulates different biological processes. Developing better tools and interventions that are effective at enhancing late-life health and longevity for both males and females depends critically on increased understanding of the biological basis of sex-specific differences in aging responses. Moreover, over the last two decades it has become evident that the sexes also respond differently to pro-longevity treatments including genetic and pharmacological interventions^10,11^. Here, we use *Caenorhabditis elegans* as a genetic model to investigate how tissue-specific degradation of the classic DAF-2 insulin-like receptor enhances health and lifespan in a sex-dependent manner.

The insulin/insulin-like growth factor 1 (IGF-1) signaling pathway (IIS) is perhaps the best described longevity-enhancing genetic pathway. Knocking out components of this pathway confers significant longevity extension in worms, flies, mice, and other animals^12^. Sex differences in the IIS pathway have also been well documented in humans and other species^13^. For example, women develop lower sensitivity to insulin than men as they age and generally have lower incidences of metabolic diseases^14^. Reduced IIS pathway activity in mice leads to a larger lifespan extension and better health outcomes in females than in males^15^. Investigating the underlying functional basis for these differences ensures that sex differences are included in the design and implementation of therapeutics aimed at enhancing health late in life.

Within *C. elegans, daf-2* encodes for an insulin-like receptor that, when mutated, can double individual lifespan^12,16,17^. This lifespan-extension phenotype is dependent on the downstream FoxO transcription factor DAF-16^16^. Since the discovery of this phenotype, the molecular components and the signaling cascade regulated by DAF-2 have been extensively studied^18^. Using the plant hormone derived auxin-induced degradation (AID) system, it has been shown that DAF-2 regulates hermaphrodite lifespan primarily through the intestine, without negatively impacting development and reproduction^2–4^. Here, we examine the sex-specific differences in longevity and healthspan in *C. elegans* by using the AID system to contrast the tissue-specific effects of DAF-2 degradation in males and hermaphrodites. Male mating represents an ideal measure of reproductive healthspan because it is a neurologically complex behavior that uses the majority of the male’s 93 sex-specific neurons to achieve^19–21^. It also declines rapidly with age^22,23^ due to behavioral and neuronal deficits^23,24^ rather than a decrease in mating drive or sperm quality. We find that ubiquitous downregulation of insulin-like signaling leads to a much larger increase in lifespan in males compared to hermaphrodites and greatly enhances male sexual function late in life. In contrast, downregulating insulin-like signaling in the male germline, unlike in hermaphrodites, decreases lifespan^3^. This work demonstrates the importance of studying sex differences in longevity and helps to expand the paradigm for investigating the sex-specific nature of these effects in one of the most important model systems for the study of the biology of aging.

### DAF-2 degradation drastically extends male lifespan

Three different groups have previously shown that ubiquitous DAF-2 degradation significantly increases hermaphrodite lifespan^2–4^. However, the effects of DAF-2 degradation on male lifespan remain unexplored. Using previously published AID strains, we asked whether the effect of ubiquitous DAF-2 degradation on lifespan is sexually dimorphic. We degraded DAF-2 in both males and hermaphrodites and measured their lifespan. We note that the whole-body DAF-2 AID strain used here produced occasional spontaneous dauers, indicative of leaky DAF-2 degradation without auxin exposure. Therefore, we used the TIR1-only strain on auxin as our negative control as reported previously^3^ (Supplemental Fig. 1). We placed 40 animals per plate (in triplicates), with hermaphrodites and males housed separately and measured their lifespan, finding that DAF-2 degradation, sex, and the interaction between the two factors have a significant effect on survival. Ubiquitous DAF-2 degradation in males led to a significant increase in survival (p < 0.0001) accompanied by a dramatic increase in median lifespan (+446%, Fig. 1a), with fully 10% of the population still alive at 106 days compared to just 15 days in controls (one way of robustly estimating “maximum” lifespan). This is much greater than the increase in hermaphrodite lifespan that we observe (+109%, Fig. 1a), which is comparable to previously reported results (+70–135%^2^, +167%^3^, and +88–117%^4^). Sex clearly plays a key role in regulating the response to IIS signaling disruption, consistent with previous findings using the canonical *daf-2* temperature-sensitive mutant^25,26^.

**Fig. 1.**
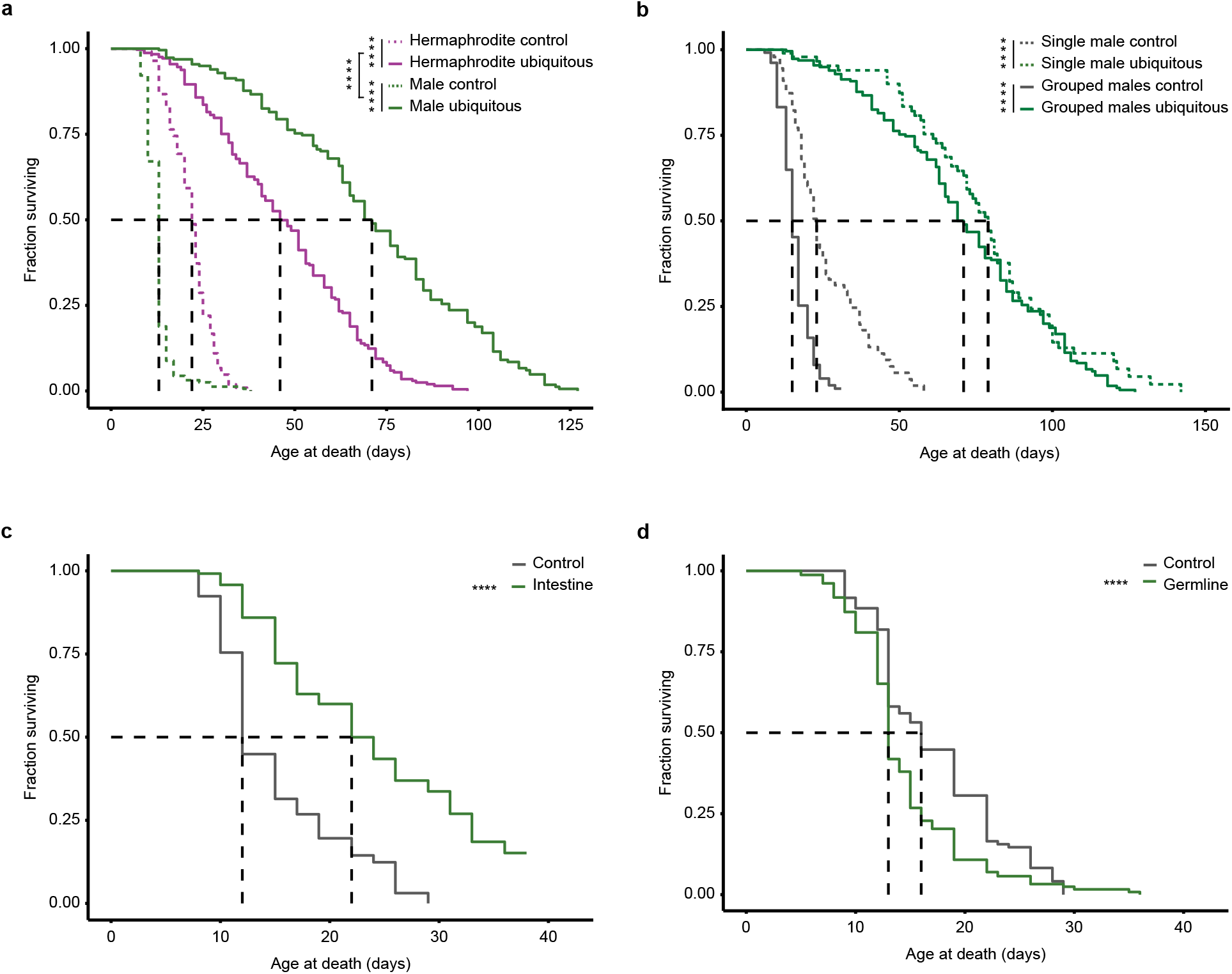
Ubiquitous and tissue-specific DAF-2 degradation extends male lifespan. (a) Kaplan-Meier curves showing survival of *C. elegans* hermaphrodites and males following ubiquitous DAF-2 degradation. Purple lines denote hermaphrodites and green lines denote males. Dashed lines denote the negative controls (TIR1-only on auxin) and solid lines denote DAF-degradation with 1mM auxin treatment. Each line represents at least two biological replicates with total n = 183–235. (b) Kaplan-Meier curves showing survival of *C. elegans* males aged on either individual or group plates. The gray line denotes the negative controls treated with ethanol and the green line denotes ubiquitous DAF-degradation with 1mM auxin treatment. The dashed lines denote single males and the solid lines denote grouped males. Each line represents three biological replicates with total n = 69–200. (c, d) Kaplan-Meier curves showing survival of *C. elegans* males following DAF-2 degradation in the (c) intestine and (d) germline. Gray lines denote the negative controls treated with ethanol and green lines denote DAF-degradation with 1mM auxin treatment. The intestine lifespan curve represents one biological replicate with n = 80–109 and the germline represents two biological replicates with n = 137–202. The black dashed lines denote the age at which 50% of the population has died. The asterisks denote *p-*values from a Cox Proportional Hazards model where ****p<.0001, ***p<.001. For additional information and the output of the CPH model, see Supplementary Table 2

Male-specific pheromones in *C. elegans* have been shown to decrease both male and hermaphrodite lifespan, resulting in “male-induced demise”^27–31^. To test for potential longevity-reducing effects of male pheromones on the above results, we individually housed males on plates and then measured the effects of DAF-2 degradation on their lifespan, comparing DAF-2 AID males on auxin to their ethanol counterparts (control). Similar to the larger scale group experiments, we found that ubiquitous DAF-2 degradation in single males also resulted in a significant increase in lifespan (+243.48%, Fig. 1b). Consistent with the overall negative effects of male-male interactions, individually reared control males lived 53.33% longer than group reared control males, although not nearly as long as DAF-2 degradation in either single or group reared males (Fig. 1b). Indeed, DAF-2 AID males display only a marginal, nonsignificant, added benefit of being housed individually vs DAF-2 AID males raised together (individual median lifespan = 79, group median lifespan = 71, p = 0.1230; Fig. 1b). This indicates that DAF-2 degradation confers close to the maximum benefit to lifespan extension in males and that eliminating male pheromones provides little additional benefit.

Next, we wanted to identify which tissues contribute to this dramatic lifespan extension. Using strains with tissue-specific promoters driving DAF-2 degradation, we targeted DAF-2 in tissues known to regulate hermaphrodite lifespan (intestine^2–4^, germline^3^, neurons ^2–4^, hypodermis^3^) and measured male lifespan. We found that intestinal degradation of DAF-2 in males extended median lifespan by 83.3% (p < 10^-14^, Fig. 1c). Germline degradation of DAF-2, however, actually decreased median lifespan by 18.8% (p < 0.0001, Fig. 1d). This negative effect of germline-specific degradation on lifespan is the opposite of what has been previously observed in hermaphrodites^3^. Further, unlike previous findings in hermaphrodites, degradation of DAF-2 in neurons or hypodermis did not alter survival (Supplementary Fig. 2a, b). Overall, we find that sex plays a significant role in modulating the magnitude of response to DAF-2 degradation in *C. elegans*, with males displaying a much greater increase in lifespan in response to this intervention than hermaphrodites.

### Ubiquitous DAF-2 degradation prolongs reproductive healthspan in old males

Lifespan and healthspan can often be decoupled, and it has become increasingly important to test the effects of pro-longevity interventions on both metrics^32–34^. We therefore tested whether the extension in male lifespan produced by DAF-2 degradation was accompanied by an extension in reproductive healthspan. We used reproductive success as our healthspan metric because it is a complex behavior that requires fidelity of multiple tissues and systems (muscles, neurons, reproductive organs)^22^, because it declines with age due to physiological changes and not because of a decrease in motivation, which is likely an important factor in reproductive behavior in hermaphrodites^23,24^, and because reproductive success is a key life history trait likely to be under strong natural selection^35,36^. We tested the mating success of young (day 1 of adulthood), middle-aged (day 7), and old (day 9) adults. These ages were selected on the scale of rapid decline of male mating success in control males (i.e., “old” males are old on the reproductive healthspan scale, even if this is close to median male lifespan). Because hermaphrodites produce sperm and can self-fertilize, we used *fog-2* pseudo-females, which are essentially spermless hermaphrodites^37^, as mating partners for the males to ensure that only cross progeny were counted. To assess mating success, one male and two virgin *fog-2* pseudo-females were allowed to mate for 24 hours, with subsequent mating success recorded via the production of any viable progeny. Using a logistic regression model, we asked whether age, DAF-2 degradation, and/or the interaction between the two factors had an effect on mating success. When DAF-2 is degraded ubiquitously, mating success at both days 7 and 9 was sustained (p = 0.057 and 0.0004, respectively), while the no-auxin controls displayed the anticipated decline in mating success. Because the males were still capable of mating at day 9, we additionally tested days 12, 15, 18, and 21 and found that ubiquitous DAF-2 degradation led to a 152.32% decrease in the overall rate of reproductive decline compared to the no-auxin control (Fig. 2a), effectively doubling the maximum age of potential reproductive success. Despite extending lifespan, intestinal DAF-2 degradation was not sufficient to preserve reproductive success in late life (Supplementary Fig. 3a). In fact, degradation of DAF-2 in any single tissue type tested here did not preserve reproductive success later in life (Supplementary Fig. 3b–d), suggesting that DAF-2 plays a multi-factorial, multi-tissue role in maintaining reproductive capacity over age.

**Fig. 2.**
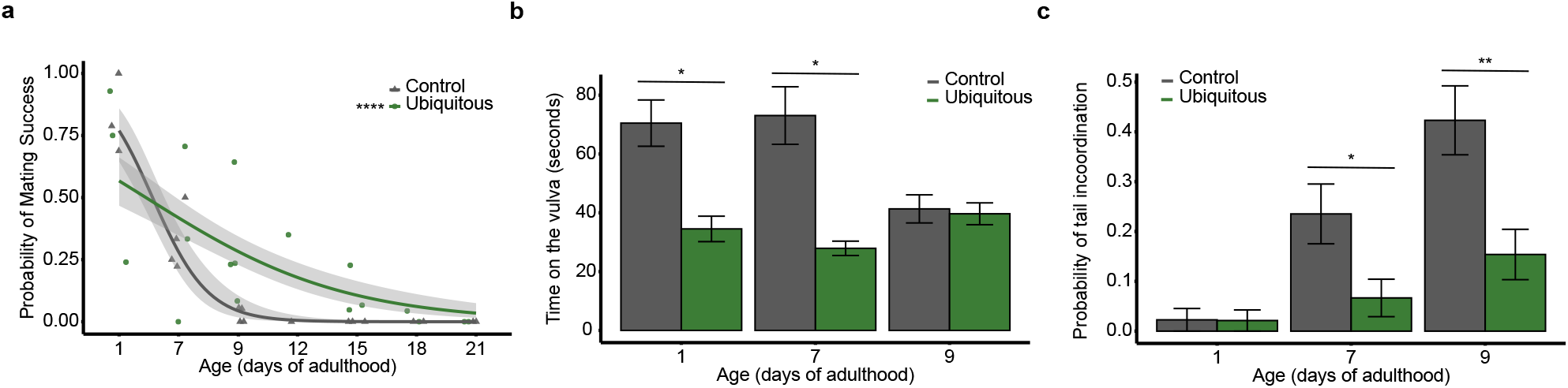
Ubiquitous DAF-2 degradation preserves late-life male reproductive success. (a) Logistic regression lines showing mating success for *C. elegans* males following ubiquitous DAF-2 degradation. The gray line denotes untreated controls, and the green line denotes DAF-2 degradation with 1mM auxin treatment. Shapes represent biological replicates, with 4–58 technical replicates in each. The gray shading around the regression lines represents SEM. A generalized linear mixed model with a binomial distribution was used to assess the effect of treatment and age (and the interaction) on mating success. (b, c) Bar graphs showing (b) the time on the vulva and (c) the probability of tail incoordination in *C. elegans* males following ubiquitous DAF-2 degradation. Gray bars denote untreated controls, and green bars denote DAF-2 degradation with 1 mM auxin. Each bar represents at least two biological replicates, with a total n = 48–52. Error bars represent SEM. A generalized linear mixed model with a binomial distribution for tail incoordination and a gaussian distribution for the time on the vulva was used to assess the effect of treatment and age (and the interaction) on behavior. The asterisks denote *p-*values the generalized linear models followed by planned comparisons where *p<.05, **p<.01, ****p<.0001. For additional information and the output of the linear models, see Supplementary Tables 3 and 4

### DAF-2 degradation lowers the incidence of male tail incoordination

For a mating event to be successful, a male must complete a series of complex behaviors. A male must locate a hermaphrodite, scan the body using its tail via backward movement, turn once it reaches the head or the tail of the hermaphrodite, locate the vulva, insert the spicule, and transfer sperm^22^. This sequence of behaviors, which is considered the most complex in *C. elegans*, declines rapidly with age^23^. To investigate which changes at the behavioral level contribute to sustained mating success, we degraded DAF-2 ubiquitously in young (day 1 of adulthood), middle-aged (day 7), and old (day 9) males and allowed them to mate with immobilized hermaphrodites then quantified different mating behaviors: turning ability, location of vulva (LOV) efficiency (vulva location success “1” or failure “0” divided by the number of passes), and the amount of time on the vulva. We note that the sample size for turning success, LOV, and time on the vulva at day 9 is limited due to many assayed males not making prolonged contact with hermaphrodites and therefore not reaching a point at which those behaviors are relevant. DAF-2 degradation in males did not have an effect on either turning ability or LOV efficiency (Supplementary Fig. 4a, b). The amount of time on the vulva, however, was significantly lower at both one and seven days of adulthood when DAF-2 was degraded (Fig. 2b). This potentially explains the lower mating output of the ubiquitous DAF-2 AID in the early ages (Fig. 2a, day 1). We also noted a novel male aging phenotype that we termed “tail incoordination” (Fig. 2c). While young males are able to perform swift sinusoidal movements (Supplementary video 1), older males seem to have a stiff tail that remains immobile while the rest of their body continues in the sinusoidal motion (Supplementary video 2). Males with this phenotype were rarely able to properly initiate the mating sequence. We found that DAF-2 degradation significantly reduces the incidence of tail incoordination in older males at both days 7 (−71.67%, p = 0.03) and 9 of adulthood (−63.64%, p = 0.003, Fig. 2c, supplementary video 3). Overall, our results suggest that sustained reproductive healthspan in older DAF-2 degraded males is driven by the decrease in the incidence of tail incoordination late in the reproductive period.

## Discussion

Sex differences in aging and the response to pro-longevity interventions have been documented widely^5,38^. Yet, the molecular mechanisms that underlie these differences remain to be fully elucidated. Here, we show that tissue-specific and ubiquitous knockdown of the insulin-like signaling pathway confer sexually dimorphic changes in longevity. We targeted the insulin-like receptor DAF-2 for degradation in the whole body of *C. elegans* males and hermaphrodites and found that the lifespan extension in males significantly exceeded that in hermaphrodites, leading to one of the largest reported lifespan extensions in *C. elegans*—or any animal—via a single intervention. Three different studies have previously shown that DAF-2 degradation in the intestine and neurons extends median lifespan in hermaphrodites^2–4^. Here, we show that DAF-2 degradation in male intestine also prolongs median lifespan dramatically, while neuronal degradation had no effect on lifespan. This difference in the effect of neuronal degradation is likely due to the sexually dimorphic nature of *C. elegans* neurons. The two sexes have several sex-specific neurons (8 in hermaphrodites and 93 in males), and some sexually dimorphic shared neurons^39^. Additionally, it has been shown recently that neuronal aging in *C. elegans* results in sexually dimorphic transcriptional changes^40^. Future work could explore whether the differences in the neuronal effect of DAF-2 degradation is via sex-specific neurons or sexually dimorphic shared neurons.

Zhang et al. also found that DAF-2 degradation in the hypodermis and germline lead to median lifespan extension in hermaphrodites^3^, while Venz et al. observed no effects of either of these tissues^2^, and Roy et al. observed no effect of germline degradation on lifespan^4^. Here, we show that hypodermis degradation had no effects on median lifespan in males while degradation in the germline actually led to a decrease in their median lifespan. Because the AID system produces a strong knockdown but not a fully null mutation, it is possible that the discrepancies among these studies is due to variable levels of DAF-2 knockdown in the hypodermis and germline. For our results, differences in development, investment, and differentiation of the germline are the most dramatic differences in morphology between males and hermaphrodites^41^. There are numerous possible explanations for the negative effects of germline-specific DAF-2 degradation on male lifespan but differentiating these will require separating potential downstream effects on primary germline tissue from the gametes themselves, which in nematodes are very substantial in size relative to the mass of the male as a whole^42^.

Incorporating both lifespan and healthspan measures into aging studies is important especially in the context of our aging population; people are living longer but suffering more diseases and disabilities in old age^43^—hardly the goal of research on aging interventions. Here, we provide additional evidence for the uncoupling of lifespan and healthspan. Despite the lifespan extension conferred by intestinal DAF-2 degradation in males, a similar effect on reproductive healthspan was not observed. In contrast, previous work had shown that in the canonical *daf-2* mutant, several health metrics such as pharyngeal pumping^44^, mobility^32,45^, learning^46^, and resistance to microbial pathogens^47^ are improved compared to wild type animals. However, in order to maintain male reproductive success into old age, multiple integrated systems and tissues need to maintain their function. Therefore, DAF-2 likely needs to be degraded in multiple tissues to produce a meaningful effect on reproductive success, which is what we observed in the ubiquitous DAF-2 AID strain. Further exploration of the effects of combined degradation in neurons and muscles, tissues known to play a role in motility and mating, on reproductive success in late life will be helpful^4^.

It has been shown previously that older males display deficits in turning behavior that is abolished in *daf-2* mutants, accompanied with improvements in the responsiveness to hermaphrodites, LOV efficiency, and turning ability^23^. We anticipated observing similar changes in the ubiquitous DAF-2 AID. However, we did not see differences in the response rate of LOV efficiency and saw a reduction in the time on the vulva. Instead, we discovered a novel tail immobility phenotype in aging males is the likely cause of sexual dysfunction, one that is ameliorated by DAF-2 degradation in the whole body. Future work could explore the basis of this phenotype and its relation to muscle health and function which had been shown previously to decline with age, leading to motor deficits that are reduced in *daf-2* mutants^4,48^, as well as other healthspan metrics such as oxidative stress resistance, motility, pharyngeal pumping, and lipofuscin accumulation^34^.

Our work shows that genetic interventions targeting the insulin-like signaling pathway extend not only male lifespan, but also their reproductive healthspan. This study expands on a framework for measuring healthspan in males as well as describing a novel tail paralysis phenotype. Taken together, these results suggest that males represent a useful tool for future aging research, as they may be more susceptible to certain interventions—fairly early in life— and provide a complex behavioral context that can be used to assess the efficacy of interventions on reproductive healthspan.

## Methods

### *C. elegans* strains and maintenance

The strains used in this study were provided by the CGC, which is funded by NIH Office of Research Infrastructure Programs (P40 OD010440). The list of strains used in this study is provided in Supplementary Table 1, many of which were sourced from Zhang et al^3^. Strains were maintained at 20°C on 60 mm plates filled with standard nematode growth medium (NGM) seeded with *Escherichia coli* OP50, unless specified otherwise. Populations were transferred every Monday and Friday to fresh plates. Plant auxin at a final concentration of 1mM for knockdown experiments was prepared by adding 400 mM auxin stock (0.350g indole-3-acetic acid added to 5mL of 190-proof ethanol) to cooled agar in a 2.5mL to 1 L ratio and poured onto plates once mixed. Plates were then covered with foil/towels and stored in light-proof boxes at 4°C after being seeded with OP50. To decrease the likelihood of spontaneous dauer formation, the ubiquitous DAF-2 AID strain was kept in a separate box from other strains, and only a few animals during maintenance were transferred to prevent overcrowding. To maintain male stocks, 8–10 males and 4–5 hermaphrodites were transferred to new plates 1–2 times a week. Stock plates older than 1–2 weeks were discarded.

### Lifespan assay

The lifespan protocol used here is based on the previously published CITP lifespan protocol^49^. A day prior to the start of the experiment, L4 males are staged visually using the male tail morphology and picked to 60 mm NGM plates seeded with *E. coli* OP50 at a concentration of 50 males per plate. Approximately 24 hours later, the adult males are transferred to experimental plates (60 mm NGM plates seeded with *E. coli* OP50, 40 males per plate, 3 plates per condition). Animals were scored three times a week and transferred to new plates weekly. Animals were scored as either alive, dead, or lost (if burrowed, missing, or walled). Mobile animals were scored as alive. Immobile animals were assayed by touching their tail or head with a platinum worm pick. If they do not respond to touch, they are scored as dead. Dead animals were removed from the plate, and live ones were counted.

### Mating assay

For each replicate, a single male was picked at the L4 larval stage and placed on its respective treatment plate: a small 35 mm NGM plate or a 35 mm auxin plate seeded with *E. coli* OP50. Each trial consisted of 16 replicates per treatment. We tested the effects of DAF-2 degradation on young, middle-age, and old males, looking at days 1, 7, and 9 of adulthood, respectively. Once the males reached their target age, they were moved to a mating plate with two 1-day old adult feminized *fog-2* hermaphrodites. These hermaphrodites do not produce sperm, therefore any progeny observed must be a product of mating with a male^37^. Animals were allowed to mate for 24 hours at 20°C before being assayed. Mating success was scored as 1 for successful mating (progeny present), and 0 for unsuccessful mating (progeny absent). Plates with contamination or missing/dead males were censored.

### Behavior assay

For each strain, 30 males were picked at their L4 stage and put on either a 100 mm NGM plate or a 100 mm auxin plate. They were then aged to their respective days of adulthood (one, seven, and nine). Following Chatterjee et al^23^, the day prior to the behavior assay, 300, L4 immobilized *unc-31* hermaphrodites were picked at their L4 stage and aged to day one of adulthood on a 100 mm NGM plate seeded with *E. coli* OP50. Five *unc-31* hermaphrodites were allowed to acclimate on a 60 mm NGM plate seeded with *E. coli* OP50 for 10 minutes. One aged male was placed on the plate and its behavior recorded for four minutes using the Dinocam software. If the male did not respond to the hermaphrodites after four minutes, it was excluded from mating measures, if it began mating in the final minute, recording time was extended to six minutes to include the entirety of the behavior. Mating was evaluated using five measurements: number of passes (the number of passes a male made over the vulva with its tail before successfully mating), LOV efficiency (Successful vulva location (1) or unsuccessful (0) divided by the number of passes over the vulva), the amount of time on the vulva (the amount of time the male kept contact with the vulva), turning success (the percentage of males that can successfully complete turning behaviors), and tail incoordination (the number of males that had upper bodies that were mobile but whose tails were immobile, over the total number of trials).

### Statistical analysis

All the statistical analyses reported here were conducted in RStudio version 4.5.1^50^. A mixed effects Cox-Proportional Hazards (CPH) model was used for each strain (single tissue degradation) to analyze survival data, where treatment (control or auxin) was used as the fixed effect and technical (Rep) and biological replicates (StartDate) were used as the random effects. To test for the effect of sex, treatment, and the interaction between the two on survival in the ubiquitous DAF-2 AID strain, a CPH model was fit where sex, treatment, and their interaction were the fixed effects, and technical (Rep) and biological replicates (StartDate) were the random effects. The coxme package version 2.2-22^51^ was used to fit the CPH model and the survival package version 3.8-3^52,53^ was used to construct the Kaplan-Meier survival curves. To analyze mating success, we fit a generalized linear model with a binomial distribution followed by paired comparisons to test for the effects of age, treatment, and their interaction on mating success within each strain. To analyze the effect of treatment on behavior, we fit a generalized linear model with a binomial distribution followed by planned comparisons to test for the effects of age, treatment, and their interaction on turning success, tail incoordination, and LOV efficiency and a gaussian distribution for the time on the vulva. The lme package version 1.1-37^54^ was used to fit the linear model and the emmeans package version 1.11.2^55^ was used for planned comparisons.

## Data availability

Raw and summary data described in this study are all publicly available on figshare (https://doi.org/10.6084/m9.figshare.30153265.v1).

## Code availability

R scripts used to analyze and visualize data in this study are all publicly available on GitHub (https://github.com/phillips-lab/DAF-2-male-manuscript.git).

## Figure legends

**Supplementary Fig. 1.**
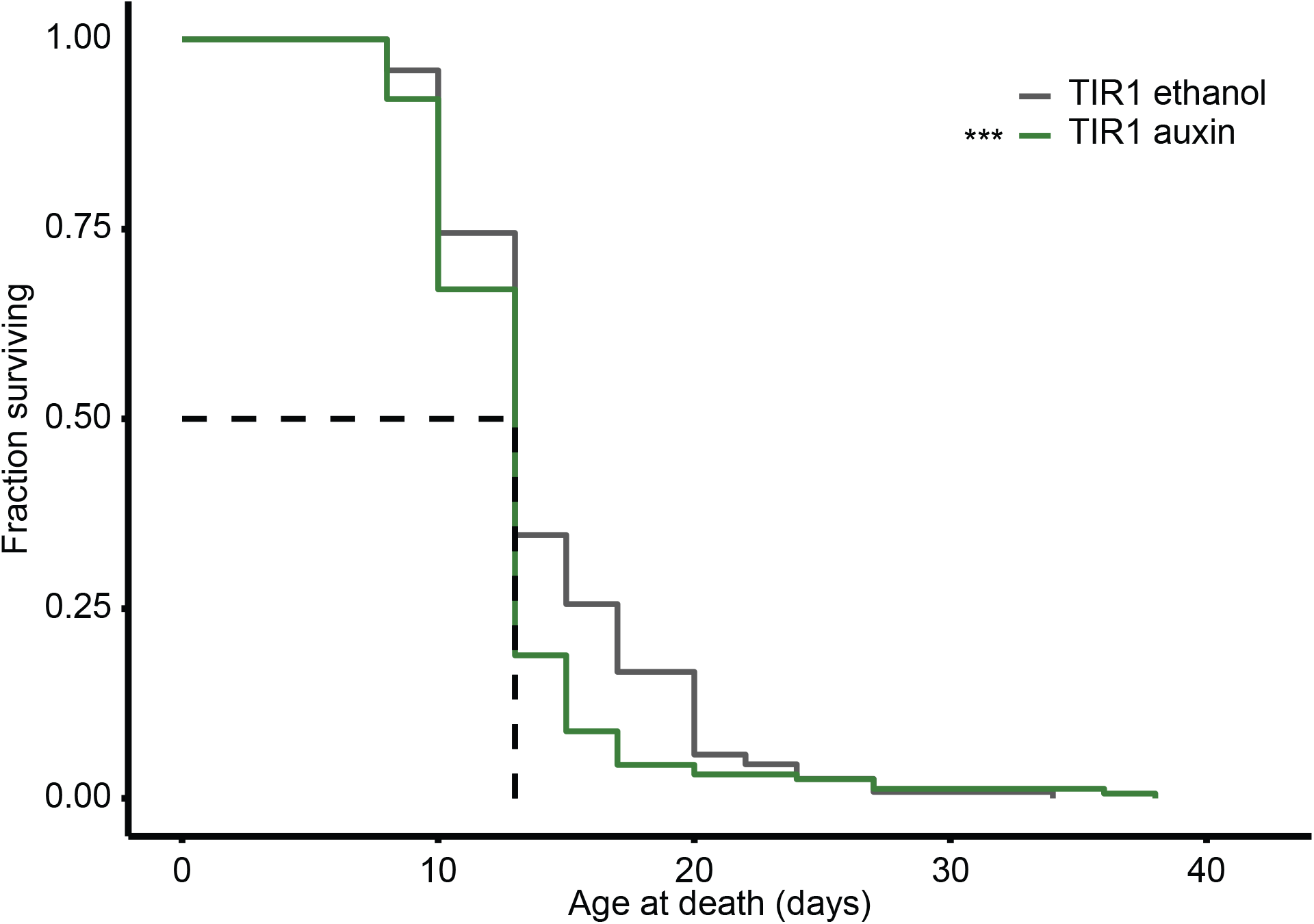
Effects of auxin treatment on male lifespan. The two major components of the AID system are: (1) a short sequence called a degron tag that is fused to the protein of interest, and (2) the plant F-box protein Transport Inhibitor Response 1 (TIR1)^56,57^. TIR1, along with other endogenous proteins, forms an E3 ubiquitin ligase complex that, when auxin is added, will target the protein of interest for degradation. However, auxin treatment alone has been previously shown to increase hermaphrodite lifespan^58^. To account for the effects of TIR1 and auxin treatment on male lifespan independent of protein degradation, we measured survival using the CA1200 strain that contains ubiquitously expressed TIR1 but no degron tag. The figure shows Kaplan-Meier survival curves for CA1200 TIR1-only *C. elegans* males. The gray line denotes the negative controls treated with ethanol and the green line denotes 1mM auxin treatment. Each line represents at least two biological replicates with total n = 179–184. Although there was a significant difference in survival between the ethanol control and auxin (p = 0.0008), there was no difference in the median lifespan for the two treatments.

**Supplementary Fig. 2.**
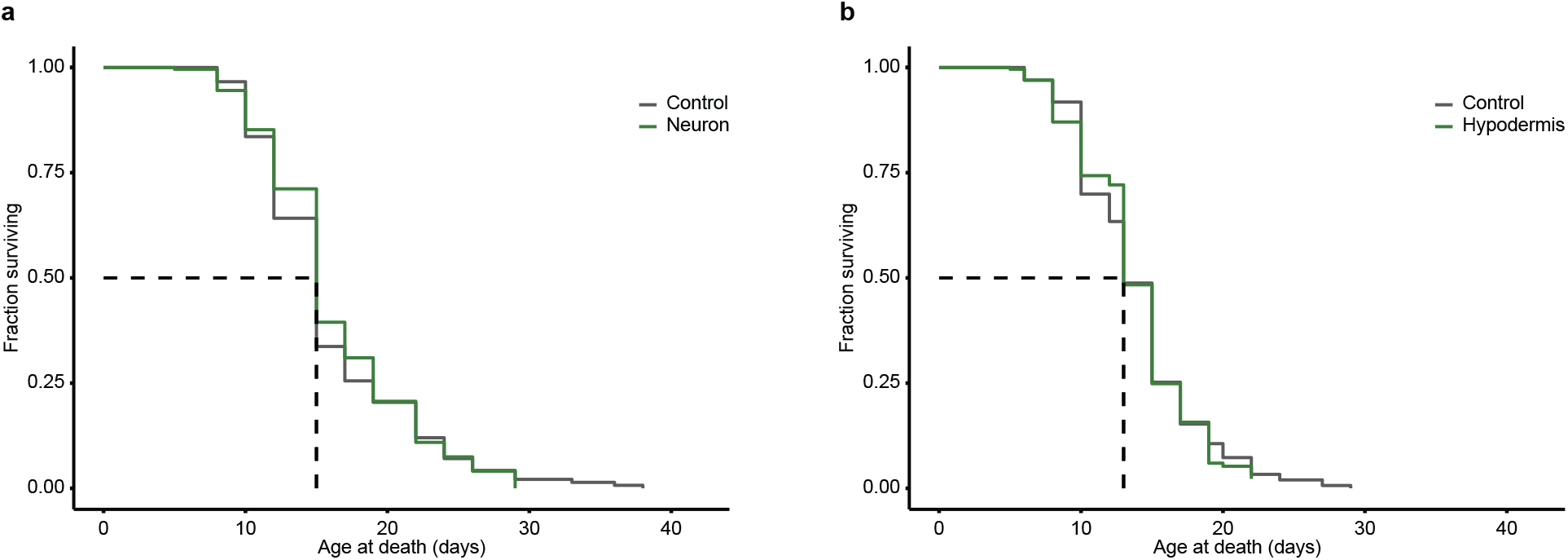
Neuron- and Hypodermis-specific DAF-2 degradation does not extend male lifespan. Kaplan-Meier curves showing survival of *C. elegans* males following DAF-2 degradation in the (a) neurons and (b) hypodermis. Gray lines denote the negative controls treated with ethanol and green lines denote DAF-degradation with 1mM auxin treatment. Each line represents at least two biological replicates with total n = 180–202.

**Supplementary Fig. 3.**
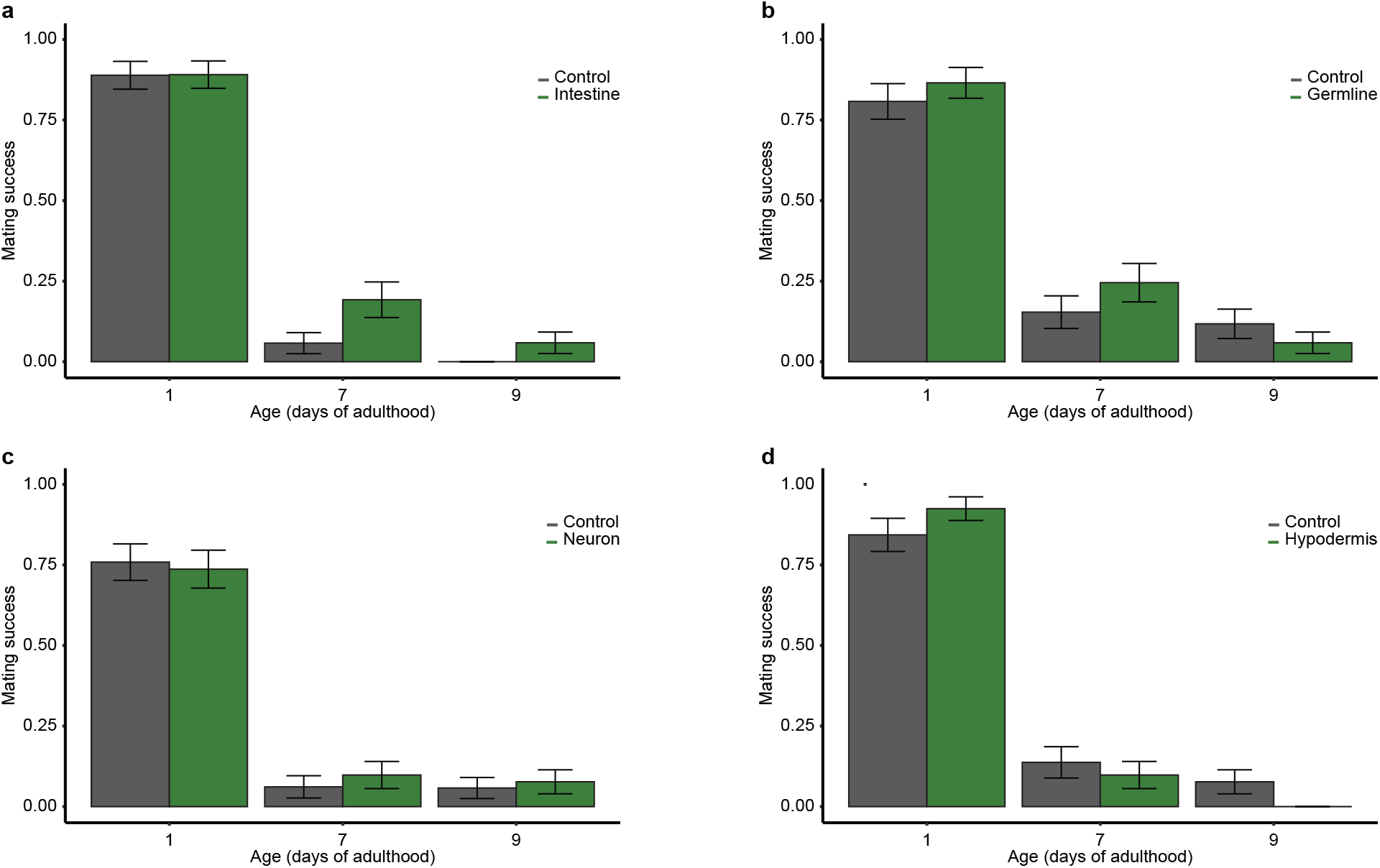
Tissue-specific DAF-2 degradation does not preserve late-life male reproductive success. Bar graphs showing mating success for *C. elegans* males following DAF-2 degradation in the (a) intestine, (b) germline, (c) neurons, and (d) hypodermis. Gray bars denote untreated controls, and green bars denote DAF-2 degradation with 1 mM auxin. Each bar represents at least two biological replicates, with total n = 49–59. Error bars represent SEM. A generalized linear mixed model with a binomial distribution was used to assess the effect of treatment and age (and the interaction) on mating success. For additional information and the output of the linear model, see Supplementary Table 3

**Supplementary Fig. 4.**
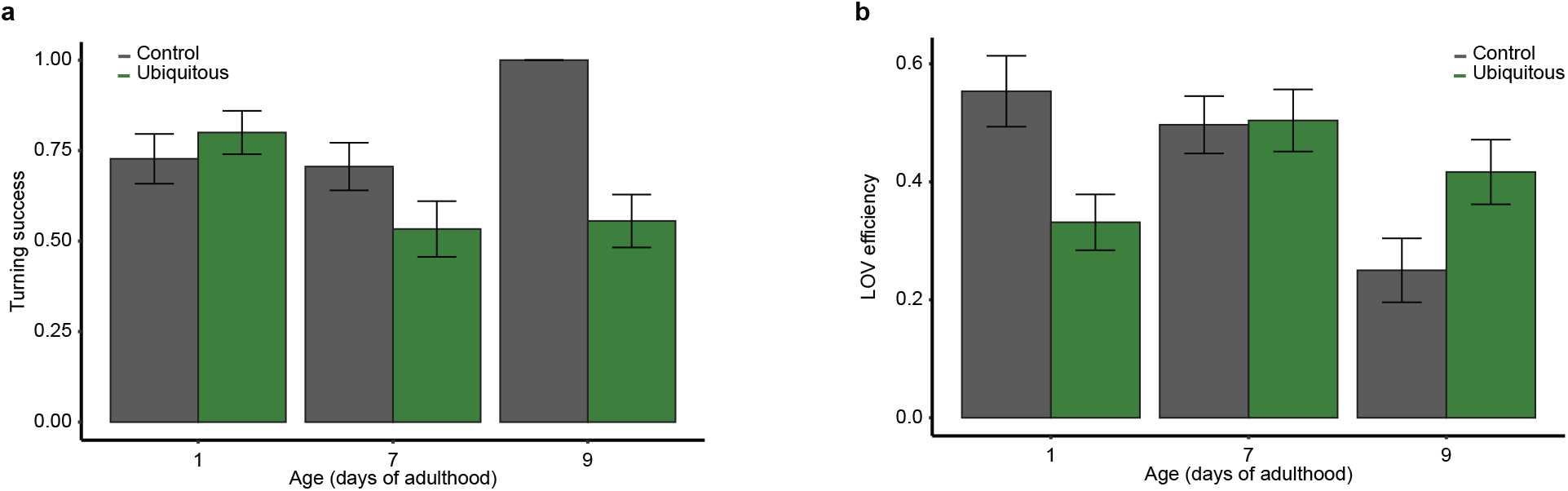
Turning success and LOV efficiency are not improved upon DAF-2 ubiquitous degradation. Bar graphs showing (a) turning success and (b) LOV efficiency for *C. elegans* males following ubiquitous DAF-2 degradation. Gray bars denote untreated controls, and green bars denote DAF-2 degradation with 1 mM auxin. Each bar represents at least two biological replicates, with total n =44–52. Error bars represent SEM. A generalized linear mixed model with a binomial distribution was used to assess the effect of treatment and age (and the interaction) on behavior. For additional information and the output of the linear model, see Supplementary Table 4

## Funding

This research was supported by grants from the National Institutes of Health to PCP (U01AG045829, U24AG056052, R01AG088629) and Training Grant Support from the National Institutes of Health to RSA (T32GM149387).

## Author information

### Contributions

RSA and HBL contributed equally to this work. RSA and PCP conceptualized the project. RSA conducted the lifespan assays, the mating assays for ages 12–21 days, conducted the statistical analysis and visualization, and co-wrote the original manuscript. HBL conducted the mating success and mating behavior assays and co-wrote the original manuscript. PCP supervised the project, reviewed and edited the manuscript, and acquired funding.

## Ethics declarations

Authors declare no competing interests.

## Notes

https://doi.org/10.6084/m9.figshare.30153265.v1

### Competing Interest Statement

The authors have declared no competing interest.

